# DNA Storage Designer: A practical and holistic design platform for storing digital information in DNA sequence

**DOI:** 10.1101/2023.07.11.548641

**Authors:** Likun Jiang, Ziyun Zou, Xinru Ruan, Xinyi Zhang, Xinyu Yu, Yinghao Lan, Xiangrong Liu

## Abstract

DNA molecules, as natural information carriers, have several benefits over conventional digital storage mediums, including high information density and long-term durability. It is expected to be a promising candidate for information storage. However, despite significant research in this field, the pace of development has been slow due to the lack of complete encoding-decoding platform and simulaton-evaluation system. And the mutation in DNA sequences during synthesis and sequencing requires multiple experiments, and wet experiments can be costly. Thus, a silicon-based simulation platform is urgently needed for promoting research. Therefore, we proposed DNA Storage Designer, the first online platform to simulate the whole process of DNA storage experiments. Our platform offers classical and novel technologies and experimental settings that simulate three key processes: encoding, error simulation, and decoding for DNA storage system. Fisrt, 8 mainstream encoding methods were embedded in the encoding process to convert files to DNA sequences. Secondly, to uncover potential mutations and sequence distribution changes in actual experiments we integrate the simulation setting for five typical experiment sub-processes (synthesis, decay, PCR, sampling, and sequencing) in the error simulation stage. Finally, the corresponding decoding process realizes the conversion of DNA sequence to binary sequence. All the above simulation processes correspond to an analysis report will provide guides for better experiment design for researchers’ convenience. In short, DNA Storage Designer is an easy-to-use and automatic web-server for simulating DNA storage experiments, which could advance the development of DNA storage-related research. And it is freely available for all users at: https://dmci.xmu.edu.cn/dna/.

**Author summary:** DNA storage technology is an emerging and promising storage technology. At the same time, DNA storage is an interdisciplinary technology that requires researchers to know both computer cryptography and biological experiments knowledge. However, DNA storage experiments are costly and lengthy, many studies have been prevented by the lack of a comprehensive design and evaluation platform to guide DNA storage experiments. Herein, we introduce DNA Storage Designer, the first integrated and practical web server for providing the simulation of the whole process of DNA storage application, from encoding, error simulation during preservation, to decoding. In the encoding process, we not only provided the coding DNA sequences but also analyzed the sequence stability. In the error simulation process, we simulated as many experimental situations as possible, such as different mutation probabilities of DNA sequences due to being stored in different bacteria hosts or different sequencing platforms. The platform provides high freedom in that users could not only encode their files and conduct the entire operation but also could upload FASTA files and only simulate the sustaining process of sequences and imitate the mutation errors together with distribution changes of sequences.

## 1. Introduction

In the era of data explosion, traditional storage methods are fast approaching a critical limit in their storage capacities (1) that are estimated to fail to satisfy global demand in 2040 (2). What’s more, the life expectancy of conventional mediums is rather short, even for magnetic tapes, which are utilized for long-term storage currently and are copied every five years for data security (3). Thus, a recent impressive development in the fields of biology and computer science involves using DNA sequences to address these issues. DNA holds an estimated information density of about 4.6×108 GB/mm3, about 6 orders of magnitude greater than the maximal density of even the most advanced magnetic tape storage system (4). Meanwhile, it could be stable for thousands of years under optimal conditions (3). High information density and long-term durability, together with other fabulous characteristics like its potentially low maintenance cost and environmental friendliness make up the expectation to let it provide wide practicality in the future. Thus, research in this area is widely carried out, and the researchers have prompted several novel technologies for the whole workflow (5-7).

Akin to classical electronic memory, a DNA-based data storage system generally involves three major steps: encoding, storage, and decoding. Unlike the binary number system of the computer, it is known-to-all that the DNA sequence has four different bases (ie. adenine, (A), thymine (T), cytosine (C), guanine(G)). Thus, the start, encoding, is to convert a binary data stream into sequences of quaternary DNA bases using a predeveloped coding schema (5). Once these DNA sequences are obtained, the next step is to synthesize them to get real DNA strands through wet experiments and store them in oligo pools. Depending on the experimental needs, the obtained DNA strands might also undergo additional steps such as storage, PCR (Polymerase Chain Reaction), sampling, and sequencing. Finally, to retrieve the data, a corresponding decoding method is required to convert information back to binary form.

Nonetheless, when it comes to actual implementation, numerous intricacies warrant careful consideration. To begin with, a primary technical obstacle pertains to the selection of an appropriate encoding method. There are many encoding methods prompted, but each way possesses unique characteristics. For example, Church et al. (7) introduced additional limitations to reduce homopolymers and repeat sequences, at the expense of lower information density, while Erlich and Zielinski (8) approach the theoretical maximum information capacity per nucleotide of DNA. What’s more, during in vitro experiments of real application, error probability changes at all stages depending on the choices and settings of experiments. As an instance, Bornhol et al. (9) reported that Illumina sequencing led to an error rate of about 1%. And Organick et al. (10) found a higher error rate of up to 10% introduced by Nanopore sequencing. It shows the truth that different choices may result in varying outcomes, making it essential to choose carefully. Apart from synthesis, each parameter chosen for each step might lead to mutations or changes in the distribution of DNA strands, which could cause information errors or even losses. Given these complexities, it is crucial to carefully consider factors such as the cost of experiments and the specificities of various documents when selecting encoding methods and experimental settings. At the same time, no simulation and evaluation platform has been reported. Due to the high cost of wet experiments, a silicon-based platform is urgently needed to assist in designing the workflow.

Herein, we introduce DNA Storage Designer, a practical and holistic web server that offers a comprehensive simulation of the entire process of DNA storage application, ranging from encoding, error simulation during preservation, to decoding. The encoding process embeds 8 mainstream encoding-decoding methods including Church’s code (7), DNA fountain code (8), Yin-yang code (11), and so forth (6, 12-14). During the error simulation stage, our server is equipped with key experimental conditions for DNA storage applications, which allows users to effortlessly configure experiments without the need for complex parameters. DNA Storage Designer grants users immense flexibility, enabling them not only to encode their files and simulate the entire process but also to upload FASTA files and solely simulate the sustaining process of sequences while mimicking the mutation errors along with distribution changes of sequences. It also gives thorough guidelines and simulated feedback based on user settings so that users could adjust their experimental plan according to the report of the website.

## 2. Design and implementation

Figure 1 presents a schematic workflow of the whole process of DNA Storage Designer. Which proposed to transfer files into DNA sequences and simulate the workflow of the whole system to guide the design of the experiment. The whole process consists of three parts, encoding, error simulation and decoding.

**Fig. 1.**
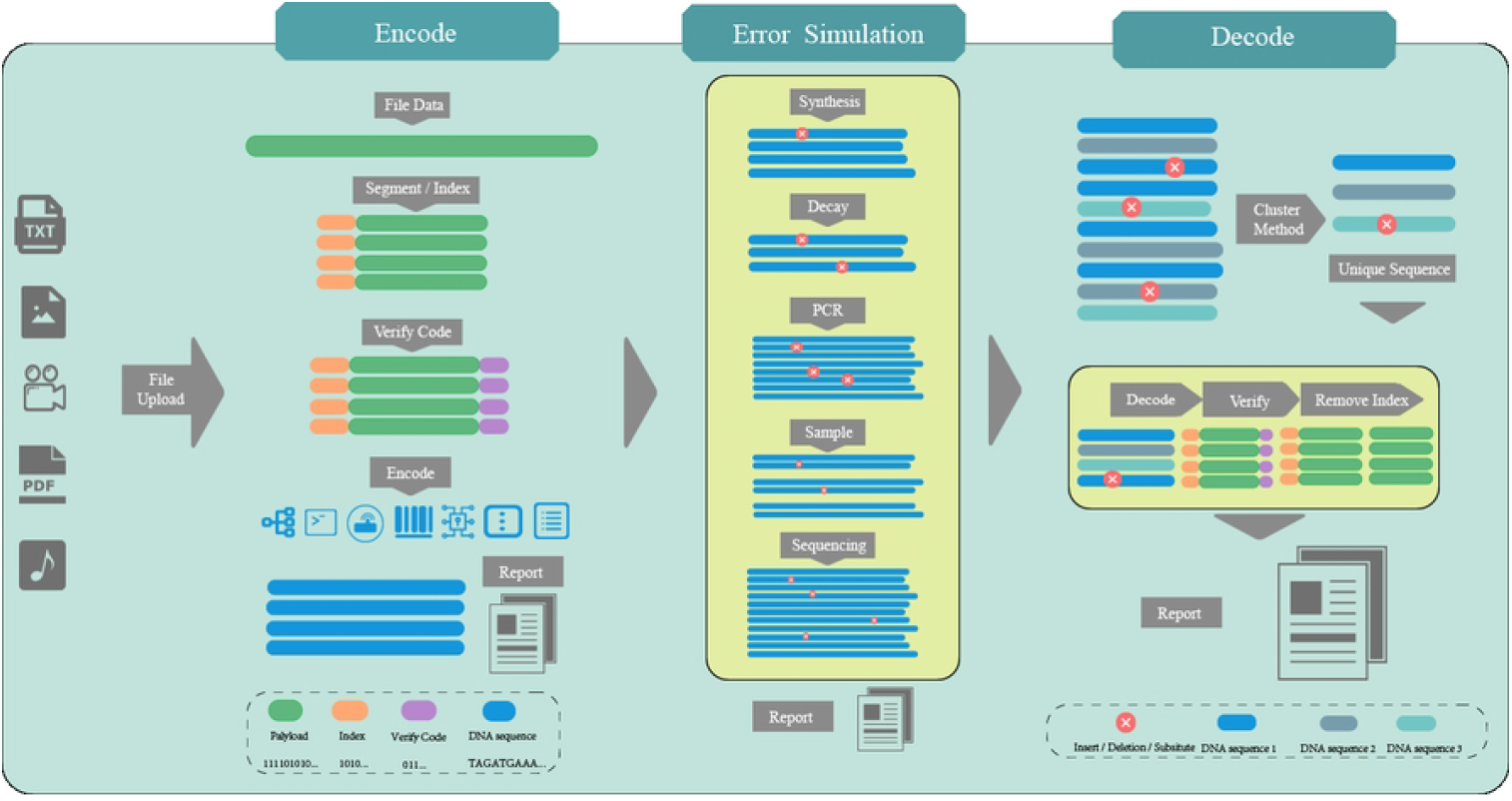
Schematic workflow of the whole process of DNA Storage Designer. First, various types of files can be uploaded to the website, the website will convert the file into binary information. After that, according to the encoding method and verify code selected by the user, each segment bit sequence will be converted into a DNA sequence after connecting the address sequence. The platform then simulated possible sequence error scenarios in the five necessary processes in DNA sequence storage experiments. Finally, the simulated sequences will be decoded to the initial digital information of the file. Each process has a corresponding analysis report.

### 2.1 Encoding

The process of encoding is exemplified in Figure 1. A digital file uploaded by users, with file types ranging from images, PDF, text, video, audio, exe, and others, is transformed into a bit matrix, where the length and encoding schema are also solicited from the users but the index length is calculated out and fixed according to the size of the uploaded file.

Initially, users should choose the encoding method to use. Currently, DNA Storage Designer provides 8 popular encoding methods, which vary from each other. Vanilla code is the most basic one, it simply transforms the data according to the naive rules: 00→A, 01 →C, 10→G, 11→T. Church’s code (7) encodes two bits per base, and Erlich (8) approach the theoretical maximum information capacity per nucleotide of DNA. Goldman et al. (12) utilize single DNA sequences to represent files with no homopolymers. Ping et al. (11) encode two binary bits into one double-stranded DNA molecule. Zan’s code (6) is proposed to only store English text, which uses a robust code book for common symbols in English. These methods hold different features, users could freely choose from based on their requirements and experiment settings. For the script of the encoding method, we refer to Chmaeleo (15), which is a robust library for DNA storage coding schemes. The details of encoding methods could be found on the “Method” page on the web server.

Then, to fix errors in the process of reading and writing DNA sequences and improve data recovery capabilities, verify code can be added optionally. Hamming code (16) and Reed-Solomon code (17) are provided. Next, users could set the segment length through the selection dot bar. The selection of fragment lengths is meticulously designed, taking into consideration a range of factors, such as file size, verification codes, and encoding methods. However, in general, it needs to meet the limitations of the current synthesis technology.

After selections are done, the file consisting of DNA segments is finally outputted and the corresponding report with the basic information of the file, together with evaluated metrics including guanine-cytosine (GC) content, repeated subsequences length, homopolymer length and the minimum free energy are directly given. Among them, GC content is a crucial indicator of DNA strand stability. It must fall within a certain interval to minimize the probability of secondary structure formation and to ensure uniform sequence coverage in the sequencing (18). Similarly, homopolymer length affects the accuracy of synthesis and sequencing (19) whereas minimum free energy could measure the quality of DNA sequences (4). Thus, we believe that these evaluation metrics could show the quality of generated sequences and provide design guidance for researchers.

### 2.2 Error simulation

The second step of the workflow is error simulation. The simulation service enables users to replicate potential errors that may arise during wet experiments, guiding designing and adjusting experiments better. It encompasses the five stages of DNA storage, synthesis, storage decay, PCR, sampling, and sequencing. As shown in Table 1, we performed full-flow simulations for these five processes while taking into account sequence errors in different experimental situations. As each stage involves high-throughput data, both in-sequences and within-sequences errors can arise, when in-sequences errors might cause information error, within-sequences errors refer to the distribution changes of sequences that could lead to information loss. As proposed and validated by Yuan et al. (20), we utilize the binomial distribution to model the sequence distribution change and within-sequence errors of each stage.

**Table 1.**
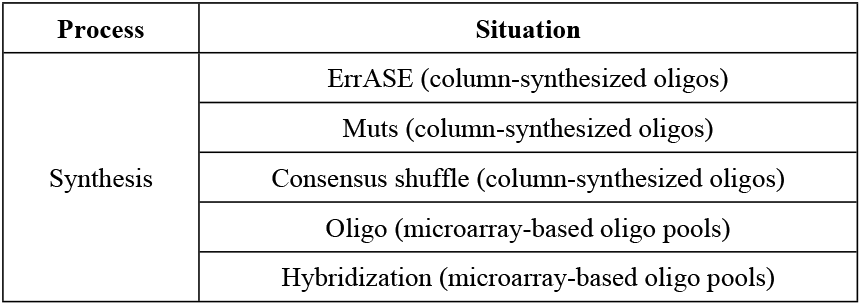
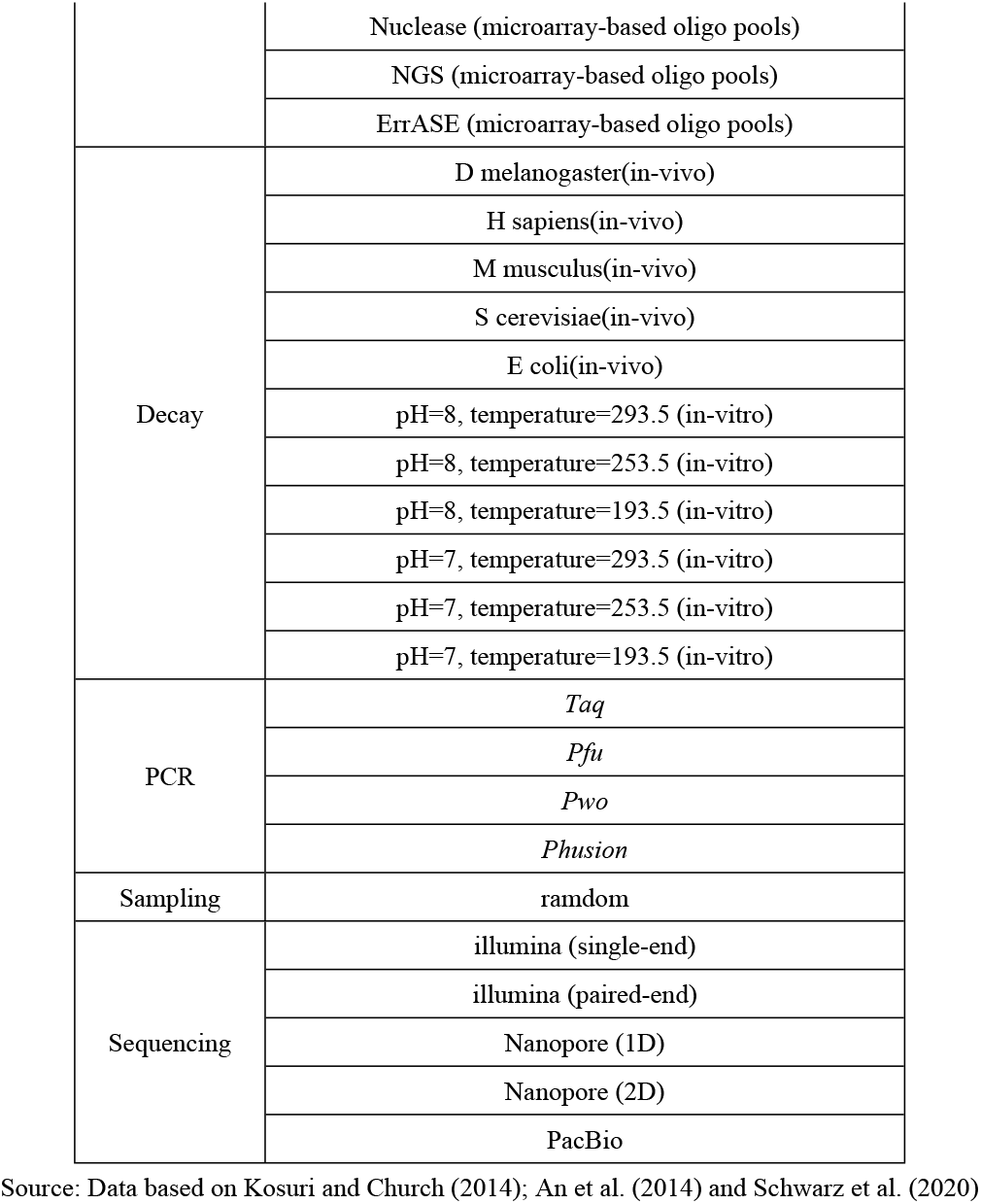
Available methods in DNA Storage Designer

During the synthesis process, some molecules might not be able to be synthesized successfully, some might be synthesized many more times than others, which causes an imbalanced sequence number distribution. What’s more, varying from different methods, the error rates and spectra are different, the available ones on our website are shown in Table 1, with published error information for different combinations of synthesis methods and error correction methods. The storage simulation simulates mutations depending on host methods and adjustable time intervals, in vitro depurination rates gained using the equation described by (21) or the Kimura model of molecular evolution (22). We provide 5 common host organisms, both Eukaryotic and Prokaryotes are considered, and 6 in-vitro experiments conditions that are commonly applied (Table 1). Also, users could simulate storage situations using the binary erasure channel or the additive white Gaussian noise channel. The polymerase used and the number of simulated PCR cycles determines the PCR error rates. We offer selections for the polymerases Taq, Pfu, Pwo and Phusion that (23) have characterized. Before sequencing, a proportion of DNA strands should be sampled from the main oligo pools, only random sequences could proceed. Thus, the sample ratio is the key parameter of this stage, and no within-sequences error will be introduced in this stage. In real experiments and applications, to read the data out, sequencing is a must. For this web server, we provide 3 kinds of prevailing sequencing platforms, Illumina (24), Nanopore and PacBio (25) with corresponding methods, in total, 6 kinds of choices. It is mentionable that substitution is the main error that occurs in this stage, especially the pair-to-pair ones, TAC-TGC and CG-CA, for example.

The report of error simulation stage consists of three parts, Steps review, Sequence distribution and Error counts. Steps review utilizes pie charts to uncover the distribution of different error types of corresponding chosen methods. During the whole process, the number of sequences, causes of errors and proportions of different types of errors change from time to time. Therefore, in Sequence distribution part, we count and compare the numbers of DNA strands with errors and the left 100% correct DNA strands for each stage using a stacked column chart, as well as show the changes in the strand numbers that contained different types of mutations using line charts. Because the effects of occurring errors are cumulative, it is reasonable that as the simulation proceeds, both the percentage of strands with errors and the average error number for all strands will increase. Thus, we count the number of strands with a different number of errors for the different stages in Error Counts section, hoping to help users to define which stage will cause most errors and might adjust their choices.

### 2.3 Decoding

The last stage is to decode the DNA sequences according to the reverse rules of the encoding ones. However, simulated DNA strands usually have a lot of redundancy, many of which have mutated errors, causing trouble with clustering and decoding. So, we embedded two clustering algorithms, CD-HIT (26) and Starcode (27), to de-redundancy and correct the data. To be specific, take CD-HIT as an example, it outputs a clustered file, documenting the sequence ’groups’ for each non-redundant sequence representative. Similarly, Starcode clustering is based on all pairs search within a specified Levenshtein distance (allowing insertions and deletions), followed by a clustering algorithm: Message Passing, Spheres or Connected Components.

Then, the clustered sequences will be decoded to obtain file binary bits (or character) information. Subsequently, the verification code and index code will be removed. Finally, we analyze the recovery information of bit fragments in the report. Similar to the report of the encoding stage, this report would also contain the basic information of users’ choices, clustering time and decoding time, as well as final results about sequence numbers and recall rate.

## 3. Usage and experiment

### 3.1 Demo Usage

To effectively showcase the capabilities of the DNA Storage Designer, we conducted a case study based on the Example file - a 140KB jpg image for Monet Claude’s Impression-Sunrise (Figure 2A, B). Upon uploading the file, we fix the segment length at 122 nt and set the hamming code as the verify code (Figure 2D). Encoded by Ping et al. (Figure 2C), the report first gives out basic file information and then evaluates several encoding results using a table with three diagrams.

**Fig. 2.**
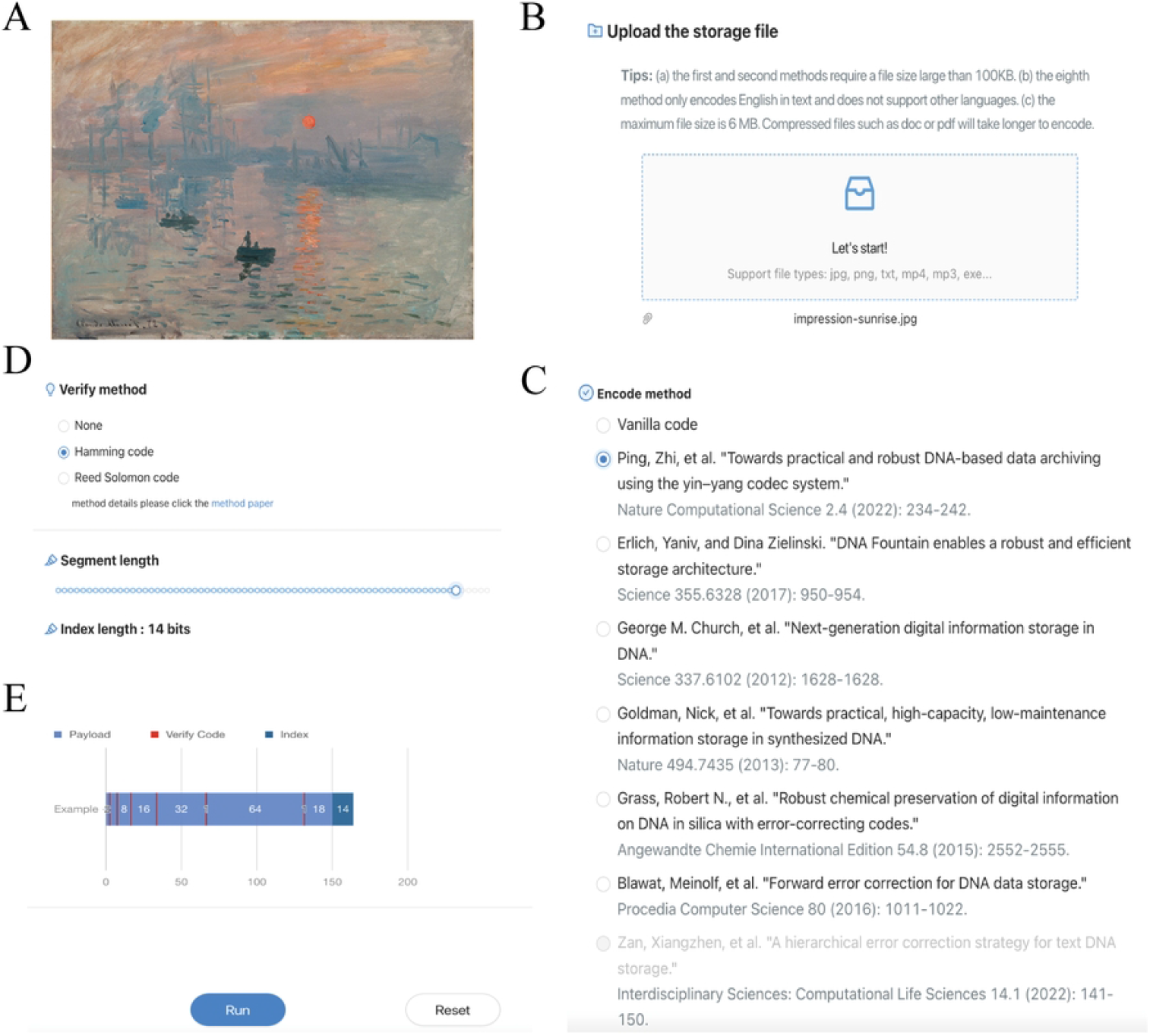
The demo run with example image file. (A) the demo file: impression-sunrise.jpg; (B) The website provides a function for users to upload files; (C) User-defined encoding method selection; (D) semi-screenshot for segment length and verify code choose which means the length of bits (01), the bits sequences will be encoded as a DNA sequence; (E) An example of a bit sequence encoded as a DNA sequence and the run button.

As shown in Table 2, the information is about how many nucleotides and sequences are generated, together with information density and so on, which first and foremost enables researchers to understand the overall situation of the encoded DNA strands. Next, we analyzed the encoded DNA sequence, as shown in Figure 3, sequence stability demonstrated by sequence minimum free energy and two diagrams display the situation of GC content, repeated sequences of randomly sampled 1000 sequences respectively. Besides, users can download the coding sequence from the report page in which the name corresponding to each sequence is the bit sequence it encodes.

**Table 2.** Sample encode information report for *impression-sunrise.jpg*.

| Name | Information |
| --- | --- |
| Encode method | Ping, Zhi, et al. |
| Segment length | 122 bits |
| Index length | 14 bits |
| Verify method | Hamming code |
| Verify code length | 8 bits |
| Encode segment length | 144 bits |
| Segment number | 8032 |
| Encoding time | 6.53 s |
| Single DNA length | 144nt |
| DNA sequence number | 8031 |
| Nucleotide counts | 1156464 *nt* |
| Information density | 0.847 *bits/nt* |
| Physical information density | 9.56E+22 *petabyte/ug* |

**Fig. 3.**
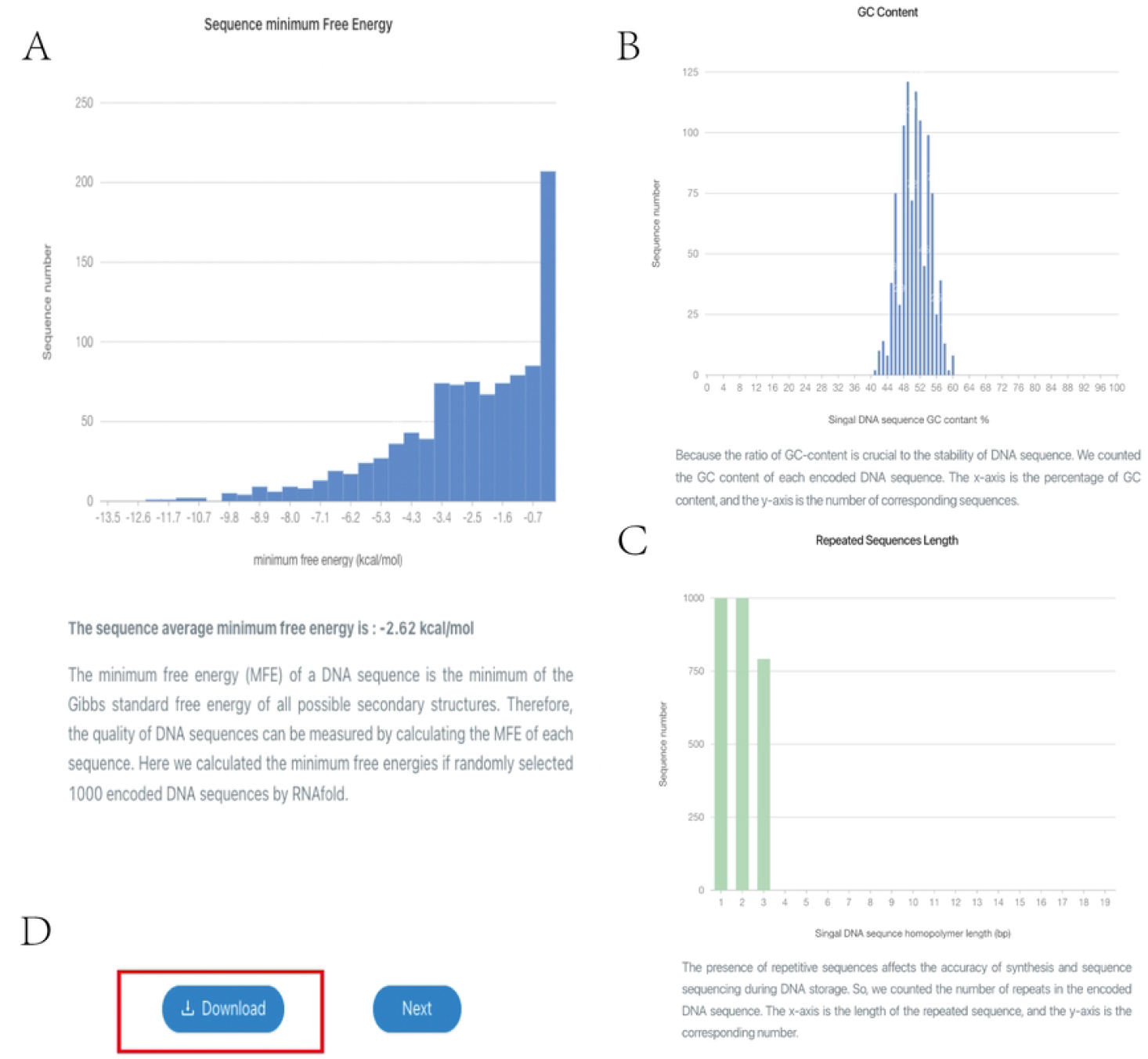
Analysis results of encoding DNA sequences for demo file. (A) The distribution of sequences minimum free energy which reflects the stability of the coding sequence; (B) Statistics on the GC content of coding DNA sequences; (C) Statistics on the repeated sequences of coding DNA sequences, each column represents the number of corresponding repeated nucleotides contained in the 1000 DNA sequence.; (D) Users can get all encoded DNA sequences by clicking the download button.

Then, we move on to the error simulation part. Under this tab, users can adjust their experimental settings on the computer, conduct simulation experiments, modify experimental parameters based on the calculated sequence analysis results, and then conduct offline experiments. To simplify, we directly press the “Default” button, which could automatically run the simulation process based on default settings for users, to conduct the demonstration. Users could also definitely adjust each parameter one by one carefully for actual usage (Supplementary Figure 1). Upon completing all selections, users can access the report page, which comprises two main sections. The simulation result provides an overview of the simulated sequence situations. During the simulation, variations in sequence distribution, density, and error occurrence are observed across different stages. To better comprehend these changes, we present “Sequence distribution” and “Error count” diagrams in Figure 4A-B, which demonstrate the change tendency of the sequences at each stage of the simulation.

**Fig. 4.**
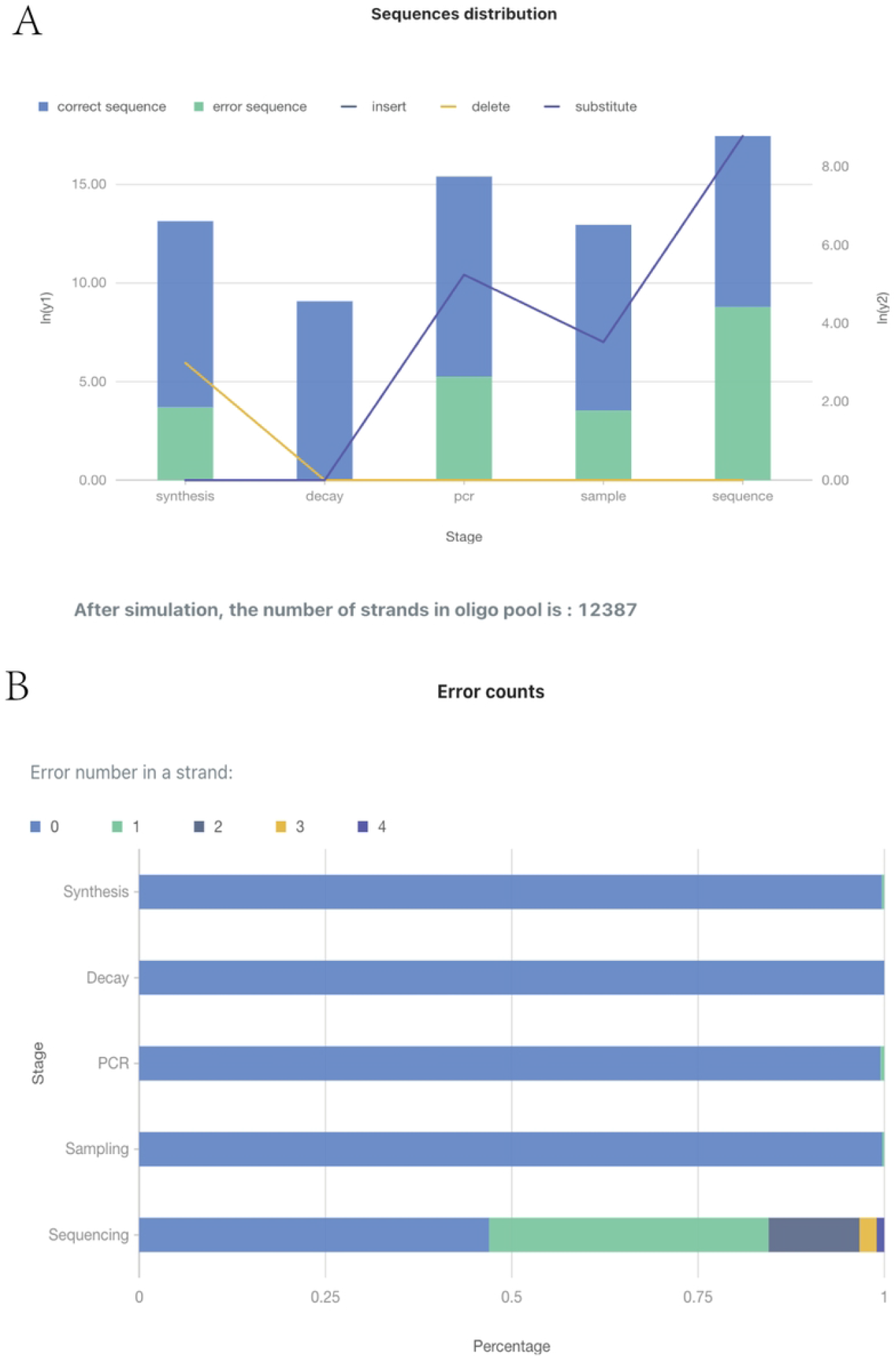
Statistical results of simulated errors in coding DNA sequences for demo file. (A) The number of correct sequences and incorrect sequences. And the number of different error types (insert/delete/substitute) in the error sequence; (B) Statistics of sequence errors for five separate experimental procedures. Users can understand the proportion of DNA strands with n errors in all strands and the changes in the proportion over time.

Finally, we use Starcode method to decode our simulated sequences above and analyze the recovery information of bit fragments in the report (Table 3). It is mentionable that the recall rate refers to the ratio of correct sequence recall ratio and the recall segment bits number stands for the ratio to the encoded counterpart respectively. These two proxies highly depend on the parameters of the simulation part. Here, because the default sampling ratio is 0.005%, and the file size is relatively small, the results could not cover all the information in the file and the recall rates are low.

**Table 3.**
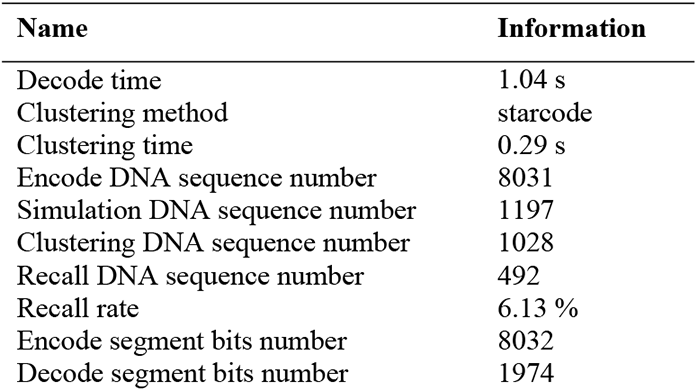
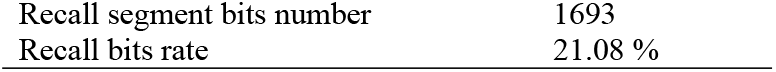
Decode information table for *Impression-Sunrise.jpg*.

| Name | Information |
| --- | --- |
| Decode time | 1.04 s |
| Clustering method | starcode |
| Clustering time | 0.29 s |
| Encode DNA sequence number | 8031 |
| Simulation DNA sequence number | 1197 |
| Clustering DNA sequence number | 1028 |
| Recall DNA sequence number | 492 |
| Recall rate | 6.13 % |
| Encode segment bits number | 8032 |
| Decode segment bits number | 1974 |
| Recall segment bits number | 1693 |
| Recall bits rate | 21.08 % |

### 3.2 Experiments and Illustration

To illustrate how our website displays variations when users select different options and how these selections impact the final results, we counted several experiments. We tested 3 files (‘impression-sunrise.jpg’,’ So far away.mp3’ and ‘Winmine.exe’) with 7 encoding methods, except Zan’s code, which is primarily intended for English texts.

The experiment results, as depicted in Figure 5, were plotted with randomly selected 1000 encoded DNA sequences from each of the three files. Compared to Vanilla code, all encoding methods exhibit improved GC content, with medians and ranges almost falling within acceptable intervals. Except for the Ping et al. (2022) code, which limits the range of GC content to 40% to 60%, the range of GC content in sequences encoded by other methods varies from file to file (Figure 5A). The minimum free energy of a DNA strand is the minimum of the Gibbs standard free energy of all feasible secondary structures. Strands with low MFE are more susceptible to secondary structures and, consequently, are more stable. While it has been reported that DNA sequences with stable secondary structures may pose challenges to sequencing or amplification during random access or backup of stored information (28-30) holds the view that a more stable strand may result in greater storage durability under appropriate conditions. Therefore, it is essential to strike a balance between stability and other factors. For the three files, Ping et al. (11), Church et al. (7), and Goldman et al. (12) encoding methods lead to lower MFE, while the others exhibit higher MFE (Figure 5B). Our website also provides users with an option to view the length of repeated sequences in encoded results. Except for Vanilla coding, all other proposed methods impose certain constraints. Among them, Goldman’s method does not allow repeated sequences, and the other three methods (George’s, Grass’s and Blawat’s) do not appear length repeat sequence greater than 3 (Figure 5C-E). However, Ping’s and Erlich’s methods will produce repeat sequences of 4 bases.

**Fig. 5.**
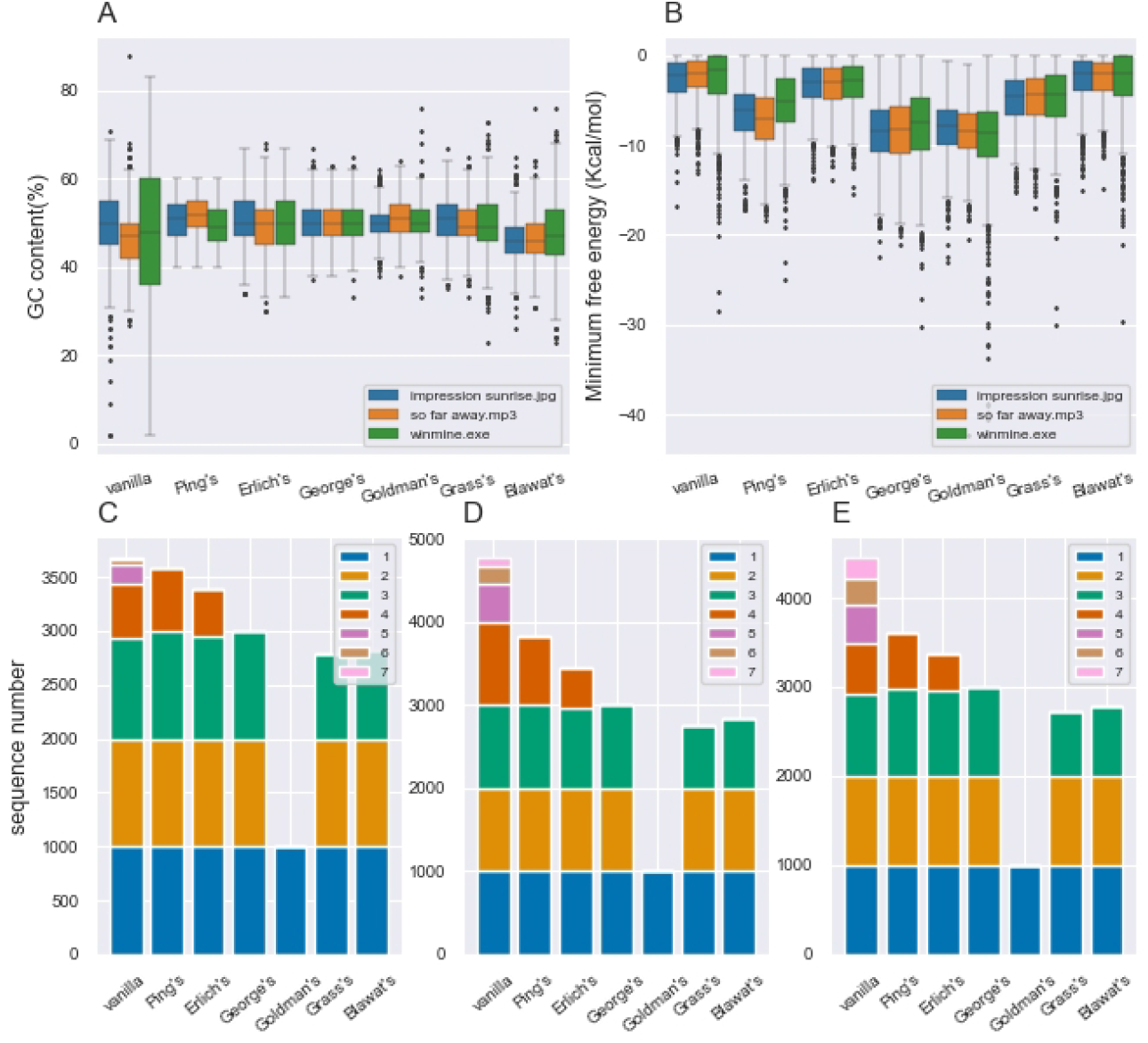
The encoding result of three files (‘impression-sunrise.jpg’,’ So far away.mp3’,’ Winmine.exe’). (A) the range of GC content of the encoded DNA sequences from the three files; (B)the distribution of minimum free energy (kcal/mol) of the encoded DNA sequences from the three files; (C) the repeat sequence number in ‘impression-sunrise.jpg’ encoded DNA sequences. Each coloured bar indicates how many repeat sequences of the corresponding length in 1000 randomly selected DNA sequences; (D) the repeat sequence number in “So far away.mp3” encoded DNA sequences; (E) the repeat sequence number in “Winmine.exe” encoded DNA sequences;

The selection of different methods, platforms, and parameters affects the result of error simulation and decoding obviously. For example, to deal with the problem we mentioned in the last part of Section 3.1, we could simply increase the sampling ratio to 100 %, and the recall rate and recall bits rate become 34.93 % and 72.22 % directly. Further, when we increase the sequencing depth from 1 to 5, the rates become 68.8% and 91.81% respectively. Also, users are encouraged to try different combinations of platforms, technologies as well as parameter settings to find the most suitable solution for their own files and application.

## 4. Conclusion

We proposed DNA Storage Designer, a practical and user-friendly web server that requires no programming knowledge. It is the first all-in-one platform to integrate the three basic processes, encoding, error simulation and decoding for DNA storage system. We embed 8 popular encoding methods that transfer digital information into DNA sequences, users could freely choose and directly transfer files into DNA sequences. The chosen encoding methods are mainly from high-impact journals such as Science, Nature, Nature Computational Science, Briefings in Bioinformatics etc., corresponding decoding processes are also included. What’s more, to simulate the real experiments of DNA storage, we also utilize the widely employed wet experiments settings in related technologies together with their error rates and spectra, to provide in-silicon evaluated results, reports and guides for experimental design. Also, the five stages, synthesis, storage, PCR, sampling and sequencing are optional. Therefore, users could decide which stages to simulate on their own. For data recovery, we utilize 2 mainstream verify codes and cluster tools to de-redundancy the sequences and help to conduct the decoding process. In general, our contribution could be summarized as:

- It is the first practical web server to simulate the whole process of DNA storage application, from encoding, and error simulation to decoding. Users could use this website to go through the whole process as well as design and modify their experiment based on the feedback.
- It incorporates 8 encoding methods together with 2 mainstream verify codes, which is currently the most inclusive one. It also has 2 cluster tools for de-redundancy purposes during decoding processes.
- It also holds a high level of usability that detailed instructions and explanations are given on the website. What’s more, each step has examples button and default settings, so users could start and use it easily.

In short, users can use our website quickly and well and are given the high degree of freedom they could upload files to go through the whole process but also can only simulate their own FASTA file or decode the files. Although DNA storage systems are not competitive for commercial use due to the limitation of current synthesis technology, it is expected that the costs for synthesis will drop significantly soon. Nevertheless, DNA storage systems allow easy and low-cost copying of media, in contrast to conventional storage systems (31). We believe that our website could provide great help for researchers and DNA storage will be implemented into daily life in the feature.

## Supporting information

**S1 Fig. Semi-screenshot of parameters for user-defined simulation error**. (A) Semi-screenshot of “choose the simulation steps”. All processes, except for synthesis, are optional and can be freely combined to customize the experiments to specific requirements. (B) semi-screenshot of choose ‘Synthesis’ parameters; (C) semi-screenshot of choose ‘Decay’ parameters; (D) semi-screenshot of choose ‘Sampling’ parameters; (E) semi-screenshot of choose ‘sequencing’ parameters;

## Acknowledgements

We thank Ping Zhi for the useful and detailed method introduction and data processing process in “Chamaeleo” approach.

